# Gestational diabetes using diverse diagnostic criteria, risk factors including dietary intakes, pregnancy outcomes and postpartum glycemic status: a nested case-control study in Ghana

**DOI:** 10.1101/582239

**Authors:** Faith Agbozo, Abdulai Abubakari, Francis Zotor, Albrecht Jahn

## Abstract

**Background:** Gestational diabetes mellitus (GDM) has risen considerably in recent years. Studies from Africa have investigated the risk factors but reported prevalence is often based on one diagnostic test/cut-off while short-term outcomes have scarcely been explored. This study estimated the prevalence of GDM using diverse diagnostic cut-offs. Associated maternal risk factors, birth outcomes and extent of attainment of euglycemia at 12 weeks postpartum were also assessed.

**Methods and Findings:** This study was an unmatched case-control nested in a prospective cohort involving 807 pregnant women recruited consecutively from five state-owned hospitals serving rural and peri-urban communities in Ghana. Dietary and obstetric risks were assessed retrospectively while physiologic measurements were repeated throughout pregnancy. Case definition was fasting venous plasma glucose (FPG) ≥5.6 mmol/l and/or single-step 75-g 2-hour oral glucose tolerance test (OGTT) ≥8.5 mmol/l measured between 20-34 gestational weeks for singleton, non-diabetic pregnant women (n=446). Participants whose random blood glucose was ≥11.1 mmol/l and glycated hemoglobin ≥6.5% were excluded. Pregnancy outcomes of 403 women were traced at delivery while 100 could be followed-up at 12 weeks postpartum. Adjusted odds ratio (aOR) for GDM was tested through unconditional logistic regression and Mantel-Haenszel statistic and the association of GDM on pregnancy outcomes was estimated by multiple logistic regression.

Prevalence per 2-h OGTT ≥8.5 mmol/l was 9.0% (n=39, 95% confidence interval [CI]; 6.3-11.6) and prevalence per FPG ≥5.6 mmol/l was 10.8% (n=49, 95% CI; 8.1-13.9); 15.9% met the case definition. Independent risk factors included excess intake of high glycemic index foods (aOR:2.91 95% CI]:1.05-8.06), obesity (aOR:2.13 CI:1.12-4.03), previous cesarean delivery (aOR:4.01 CI:1.08-14.76) and antenatal care in a primary facility (aOR:4.951 CI:1.87-3.76). A unit rise in blood glucose significantly increased maternal blood loss and birthweight. Adjusting for covariates, adverse birth outcomes were perineal tear (Aor:2.91 CI:1.08-5.57) and birth asphyxia (aOR:3.24 CI:1.01-10.44). Cesarean section (aOR:1.9 CI:0.97-3.68), large for gestational age (aOR:2.7 CI:0.86-5.05) and newborn resuscitation (aOR:2.91 CI 0.94-9.01) were significant at 10%. At 12 weeks postpartum, 30% of the GDM cases were unable to achieve euglycemia. Different estimates could be obtained if other diagnostic criteria were used.

**Conclusions:** Findings show an increasing prevalence of GDM in peri-urban and rural settings highlighting the need to strengthen primary facilities to test and refer cases for management. Diet and adiposity are key risk factors necessitating lifestyle modification interventions focusing on nutrition education and weight control. GDM-exposed newborn need close monitoring as birth asphyxia which is a key outcome is likely to compromise neonatal survival. Postpartum follow-up of cases is crucial to avert transition of GDM into active diabetes.

## Introduction

Gestational diabetes mellitus (GDM) is glucose intolerance that affects 1–14% of all pregnancies globally [1]. In 2017, 16.2% of live births experienced hyperglycemia in pregnancy, of which 86.4% was accountable to diabetes first recognized in pregnancy [2]. Predisposing factors like ethnicity, family history of diabetes, advanced maternal age, short stature and previous bad obstetric history are non-modifiable. Focusing on modifiable risk factors like obesity, high parity, gestational weight gain, sedentary lifestyle, excess caloric intake, abnormal lipid profile and hypertension could reduce the prevalence [3–7]. With a current GDM prevalence of 14% in sub-Saharan Africa [4], Ghana like many developing countries is struggling to tackle the nutrition and epidemiological transition. The double burden of undernutrition and obesity resulting from unhealthy lifestyle, food insecurity, sedentary behaviors and urbanization is promoting insulin resistance [7].

Adverse outcomes linked to GDM include preeclampsia, postpartum hemorrhage, obstructed labor, cesarean delivery, macrosomia, birth trauma, asphyxia, neonatal hypoglycemia and perinatal mortality [5, 8, 9]. GDM affects breastfeeding [7] and long-term cardio-metabolism of mother-offspring pairs [6, 7, 10]. Despite recommendation by the International Association of Diabetes and Pregnancy Study Group (IADPSG) on ‘one-step’ testing of all pregnant women using fasting glucose and the ‘gold standard’ 2-hour oral glucose tolerance test (OGTT) [11], disagreements exist on diagnostic thresholds [12]. It is argued that lower diagnostic thresholds would increase GDM prevalence and burden health systems in low-resource settings which might not be prepared to manage the sheer number of cases [13].

While obesity and positive family history of diabetes are known risk factors, in rural areas in Ghana, maternal obesity (4.3%) [14] and type 2 diabetes (4.0%) [15] are low. Studies on GDM prevalence and the risk factors are few in Ghana and adverse pregnancy outcomes are not well established. Also, population-based prevalence from peri-urban and rural settings is unavailable. Current data on GDM in Ghana shows a prevalence of 9.3% using 2-h OGTT >8.5 mmol/l [16]. Given that this study was conducted in the largest referral/teaching hospital in Ghana where 92.5% of the study participants were urban dwellers, findings might not be representative of the typical Ghanaian population where lower population-based prevalence is hypothesized.

In Africa, studies have assessed risk factors, but literature on short-term outcomes is limited. Therefore, we prospectively followed a cohort of pregnant women and obtained data on GDM prevalence from lower levels of healthcare delivery. Socio-demographic, dietary, anthropometric, obstetric and physiologic factors that increased risk for GDM were assessed as well as the associated short-term birth outcomes and maternal postpartum glycemic status.

## Materials and Methods

### Design

This observational study was an unmatched case-control nested in a prospective cohort study in which the primary aim was to validate accuracy of screening tools for GDM and assess associated birth outcomes. Synopsis of the methods applied is shown in Fig 1. Participants were tested for GDM in the second to third trimesters of pregnancy, birth outcomes assessed at delivery and glycemic status monitored at 12 weeks postpartum. Physiologic measurements were taken at every antenatal care (ANC) visit while birth outcomes were extracted from the facilities’ delivery records. The case-control component involved obtaining data on socio-demographic characteristics, measuring weight, height, random blood glucose and glycated hemoglobin and retrospectively assessing dietary intake, medical and obstetric risk factors for GDM.

**Fig 1.**
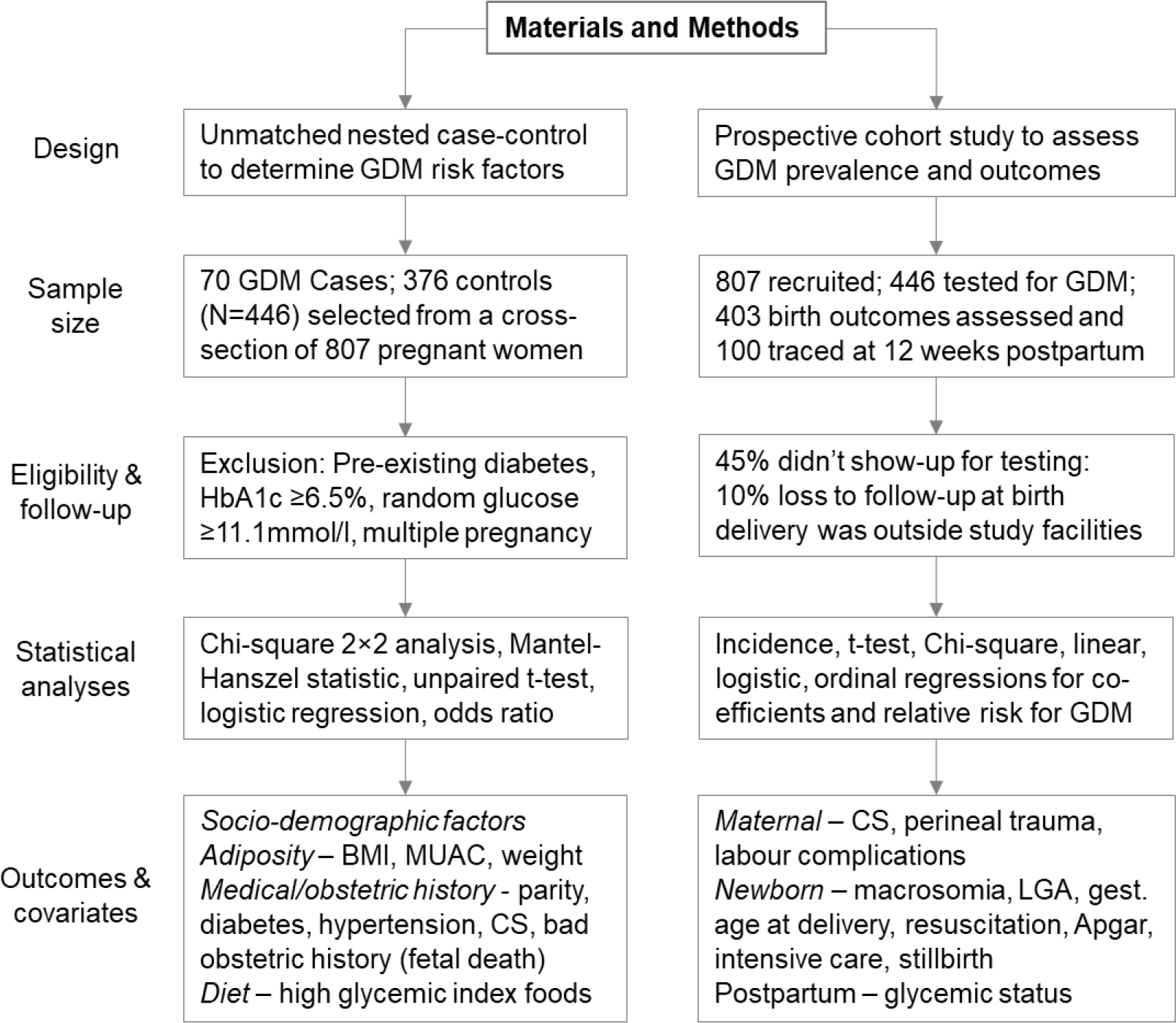
Synopsis of the methodology.

### Setting

Generally, in Ghana, ANC and delivery services are provided at the secondary level and is administrated through the Ghana Health Service. Hence participants were proportionately recruited from one primary hospital (15%), three secondary hospital (70%), and one regional-level referral hospital (15%) serving rural and peri-urban communities in the Volta region, Ghana. The primary facility provides basic emergency obstetric and newborn care (EmONC) while the secondary facilities provide comprehensive EmONC signal functions. Women in their reproductive ages (15-49 years) constitute 46.8% of the 1,098,854 female population in the region. Total fertility rate for women aged 15-49 years is 3.2 per woman compared to the national average of 3.3 while the completed family size for women age 12-54 years is 5.1 per woman [17]. Fertility rates of rural dwellers is relatively higher but there is little difference in teenage fertility in urban and rural areas.

### Sample size

The sampling frame was 3,093 pregnant women who had registered for ANC in the five study facilities at the time of conducting the study from June 2016 to April 2017. The estimated sample size was 416 and has been justified elsewhere [14]. This number (n=416) was increased to 807 to account for participant attrition. The eligibility criteria were singleton pregnant women without pre-existing diabetes who registered for ANC in the first trimester of pregnancy and intended to deliver in the study facilities. Participants tested for random blood glucose and glycated hemoglobin (Hb1Ac). Values ≥11.1 mmol/l and ≥6.5% (7.8 mmol/l) respectively were suggestive of pre-existing diabetes and such cases were excluded from the study. Overall, 807 participants were booked for GDM testing of which 490 representing 55% reported but 44 were excluded due to non-fasting. Thus 446 participants tested for GDM; 403 were traced at delivery while 100 were followed-up at 12 weeks postpartum.

### Data collection tools and procedures

#### Glycemic status

Testing was done between 20-34 gestational weeks using fasting venous plasma glucose (FPG) following 12 hours overnight fast. Seventy-five-gram anhydrous glucose was dissolved in 300 ml of water and drank under direct observation after which the 2-h postprandial glucose was measured. In addition, fasting lipid profile was checked. Diagnosis of GDM from FPG was in line with the NICE guidelines (National Institute for Health and Care Excellence) [18] whereas diagnosis of GDM from 2-h OGTT was in line with the IADPSG/WHO guidelines [11, 13] (Table 1). Basis for choosing these two thresholds emanate from a previous study where FPG ≥5.6 mmol/l and 2-h OGTT ≥8.5 mmol/l were found to be more diagnostically accurate and had higher disease prediction [14]. Also, lowering of thresholds for GDM diagnosis have spark concerns of overdiagnosis [19], hence the decision to restrict to using higher diagnostic thresholds for both FPG (≥5.6 mmol/l) and 2-h OGTT (≥8.5 mmol). Since only one abnormal value is needed for diagnosis [13], the case definition and diagnosis of GDM was based on fasting plasma glucose value ≥5.6 mmol/l and/or 2-h OGTT value ≥8.5 mmol/l.

**Table 1.** Prevalence of gestational diabetes according to common diagnostic criteria

| Diagnostic criteria | Fasting glucose (N=446) |  | 2-h OGTT (N=435) |  | Fasting and/or 2-h OGTT positive<sup>d</sup> |
| --- | --- | --- | --- | --- | --- |
|  | Cut-off mmol/l | Prevalence % | Cut-off mmol/l | Prevalence % | Prevalence % |
| IADPSG<sup>a</sup>/WHO/FIGO/ADA | 5.1 | 23.8 | 8.5 | 9.0 | 26.5 |
| NICE | 5.6 | 10.8 | 7.8<sup>c</sup> | 14.3 | 20.3 |
| CDA | 5.3 | 16.9 | 9.0 | 5.1 | 18.9 |
| This present study<sup>b</sup> | 5.6 | 10.8 | 8.5 | 9.0 | 15.9 |

IADPSG: International Association of Diabetes in Pregnancy Study Groups; WHO: World Health Organization; FIGO: International Federation of Gynecology and Obstetrics; ADA: American Diabetes Association; NICE: National Institute for Health and Care Excellence; CDA: Canadian Diabetes Association.
<sup>a</sup> IADPSG cut-off adopted by the Australian Diabetes in Pregnancy Society and Brazilian Society of Diabetes.
<sup>b</sup> This study used the NICE cut-off for fasting plasma glucose ( $\geq 5.6$ mmol/l) and the WHO cut-off for 2-h OGTT ( $\geq 8.5$ mmol/l)
<sup>c</sup> NICE cut-off for 2-h OGTT is used by the Diabetes in Pregnancy Study group in India but the Indian group test for 2-h OGTT irrespective of the fasting state of the woman
<sup>d</sup> GDM is diagnosed made when one or both of the diagnostic cut-off values is exceeded

#### Exposure factors

Face-to-face interviews were conducted with 807 participants to obtain information on socio-demographic characteristics (e.g. marital status, educational level of the couple, place of residence), anthropometric indices, obstetric history and dietary intakes. Weight and height were measured in the first trimester to estimate the body mass index (BMI). In accordance with recommendations from the Institute of Medicine on ideal pregnancy weight gain based on a woman’s BMI group, a high GDM risk was considered if the pregnancy weight gain was above the threshold for her BMI group [20]. The BMI groups and corresponding pregnancy weight gains are as follows: underweight (<18.5), 12.5-18.0 kg; normal weight (18.5-24.9), 11.5-16.0 kg; overweight (25.0-29.9), 7.0-11.5 kg; and obese (≥30), 5.0-9.0 kg. Mid-upper arm circumference (MUAC) was measured in each of the trimesters of pregnancy while pregnancy weight change was monitored very month to assess adiposity. MUAC cut-off was based on the population’s median value. Obstetric history obtained included gravidity, parity, spontaneous abortions and previous cesarean delivery, macrosomia and stillbirth. Family history of diabetes and hypertension were evaluated while physical assessments (e.g. glycosuria, proteinuria, blood pressure) were taken at every antenatal care visit. Dietary intake was assessed using a 24-hour recall to obtain qualitative data on intake of high glycemic index (GI) foods including snacks and beverages during the day prior to the interview. Each carbohydrate-containing food consumed that contributed >70% of the GI value (e.g. white bread, polished rice, processed cassava and corn meals, ripe plantain, table sugar, pasta, pineapple, watermelons, soda drinks) was assigned a score of one. Cumulative scores were classified into adequate (1-4 high GI foods per day); moderate (3-4 intake of high GI foods per day) and excess (≥5 high GI foods per day) intake of high glycemic index foods.

#### Outcome measures

Primary maternal outcomes were cesarean delivery and perineal trauma while the secondary outcomes were preeclampsia (defined as concurrent high blood pressure and proteinuria with or without edema) and post-partum hemorrhage (estimated blood loss >500ml). Primary newborn outcome was adiposity and survival. Adiposity was assessed using three indicators: (1) macrosomia defined as birth weight ≥4 kg regardless of gestational age at birth; (2) large for gestational age (LGA) defined as birth weight >90^th^ percentile per the InterGrowth study standards accounting for gestational age at birth and sex of the newborn; and (3) Ponderal Index (PI) defined as newborn weight (g)/length (cm^3^) ×100 and classified as small for gestational age (<2.0), marginal (2.0-2.5), normal (2.5-3.0.) and large for gestational age (≥3.0). Newborn survival was assessed using four indicators: (1) Apgar score at one and five minutes, (2) newborn resuscitation, (3) admission to neonatal intensive care unit (NICU) and (4) perinatal death. Secondary newborn outcomes were gestational age at birth and capillary heel prick blood taken between 1-2 hours after birth to assess for hypoglycemia.

### Statistical analysis

Descriptive and inferential analyses were run in Stata software (version 14.2). Scale variables were analyzed using unpaired t-test and reported as means with the standard deviations (SD) while categorical variables were analyzed by Chi-square test and reported as frequency distributions. P-values <0.05 and confidence intervals (CI) excluding one indicated statistical significance. The case-control component was analyzed using a dichotomous outcome (GDM present or absent) tabulated in a two-by-two table with dichotomous risk factors. Aided by the Cochran-Mantel-Haenszel statistic, unconditional logistic regression model was fitted to generate the crude estimates of association between exposures and GDM outcome. Confounding variables were adjusted for by stratifying the exposure variables into sub-groups and computing the multivariate binary logistic regression to obtain the adjusted odds ratios (aOR). Concerning the pregnancy outcomes, simple linear regression analysis was modelled to estimate the strength and direction of association for a unit rise in FPG and 2-h OGTT on each pregnancy outcome. A correlation matrix was computed to identify collinearity and possible confounders. Then, a multivariate binary logistic regression model was run to estimate the adjusted association of GDM on pregnancy outcomes. Interaction terms were considered in the final model selection.

### Ethical approval and consent

The study was approved by the Ghana Health Service Ethics Review Committee (GHS-ERC-GM 04/02/16) and the Institutional Review Board of Heidelberg University Medical Faculty (S-042/2016). Voluntary written informed consent was obtained from all study participants including pregnant women less than 18 years who were ethically considered as emancipated adults.

## Results

### Prevalence of gestational diabetes

Fasting plasma glucose of 446 participants and 2-h OGTT of 435 participants were obtained. Prevalence of GDM per 2-h OGTT ≥8.5 mmol/l was 9.0% (n=39, 95% CI; 6.3-11.6) and prevalence per FPG ≥5.6 mmol/l was 10.8% (n=49, 95% CI; 8.1-13.9). Participants who met the case definition of 2-h OGTT ≥8.5 mmol/l and/or FPG ≥5.6 mmol/l were 15.9% (n=70/446; 95% CI; 12.5-19.3). Only 3.9% (n=17/433; 95% CI; 2.1-5.8) were positive for both 2-h OGTT and FPG. Presented in Table 1 is GDM prevalence according to the common diagnostic criteria.

### Maternal health characteristics

Seventy participants met the GDM case definition and the remaining 376 served as controls, representing one case to about approximately five controls. Mean maternal age was 28.44 years (SD=6.13); the minimum and maximum ages were 15 and 54 years respectively. Five (1.8%) participants were between 15-17 years while only one participant was above 49 years. A third (n=153) were primiparous mothers. The cases were significantly older, had higher adiposity, gravidity and parity (Table 2).

**Table 2.** Unpaired t-test comparing characteristics of cases and controls for continuous variables

| Continuous variables | Mean (standard deviation) |  |  | *P-value* |
| --- | --- | --- | --- | --- |
|  | Total (N=446) | GDM<sup>a</sup> (n=70) | No GDM (n=376) |  |
| Fasting glucose<sup>a</sup> | 4.56 (0.99) | 5.70 (0.79) | 4.28 (0.69) | -- |
| 2-h OGTT <sup>a</sup> | 6.33 (1.72) | 7.82 (1.73) | 5.90 (1.14) | -- |
| Maternal age (years) | 28.44 (6.13) | 29.82 (6.80) | 28.19 (5.99) | 0.036 |
| Gravida | 2.73 (1.49) | 3.17 (1.57) | 2.66 (1.47) | 0.010 |
| Parity (live children) | 1.35 (1.31) | 1.75 (1.45) | 1.28 (1.27) | 0.007 |
| Weight (kg) | 62.98 (13.06) | 66.38 (14.32) | 62.37 (12.75) | 0.020 |
| Height (cm) | 162.36 (9.33) | 162.93 (7.94) | 162.26 (9.54) | 0.604 |
| BMI (kg/m<sup>2</sup>) | 23.48 (4.43) | 24.77 (4.82) | 23.21 (4.30) | 0.026 |
| MUAC (cm) | 28.33 (3.69) | 29.51 (4.40) | 28.08 (3.48) | 0.016 |
| Weight change<sup>b</sup> | 11.16 (5.18) | 11.67 (5.21) | 11.06 (5.18) | 0.445 |
| Systolic BP (mmHg) | 108.06 (11.49) | 109.88 (11.91) | 107.74 (11.40) | 0.154 |
| Diastolic BP (mmHg) | 65.39 (9.37) | 65.58 (10.23) | 65.36 (9.22) | 0.859 |
| Total cholesterol (mmol/l) | 5.50 (1.28) | 5.32 (1.37) | 5.54 (1.27) | 0.189 |
| Triglycerides (mmol/l) | 3.05 (1.65) | 2.88 (1.23) | 3.08 (1.71) | 0.331 |
| HDL cholesterol (mmol/l) | 1.47 (0.44) | 1.49 (0.49) | 1.47 (0.43) | 0.596 |
| LDL cholesterol (mmol/l) | 2.66 (1.07) | 2.51 (1.04) | 2.68 (1.08) | 0.201 |
| VLDL cholesterol (mg/dl) | 1.35 (0.76) | 1.30 (0.60) | 1.36 (0.78) | 0.542 |
| Coronary risk | 4.14 (2.37) | 3.80 (1.20) | 4.20 (2.52) | 0.180 |
| Intake of high GI foods<sup>c</sup> | 2.62 (1.14) | 2.87 (1.24) | 2.56 (1.12) | 0.242 |

BMI, body mass index; MUAC, mid-upper arm circumference; BP, blood pressure; HDL, high density lipoprotein; VLDL, very low-density lipoprotein; GI, glycemic index.
Apart from the lipid profile, all other assessments were taken in the first trimester.
<sup>a</sup> The case definition is 2-hour OGTT $\geq 8.5$ mmol /l and/or the fasting plasma glucose $\geq 5.6$ mmol/l.
<sup>b</sup> Pregnancy weight change was estimated by subtracting weight measurement at delivery from the weight in the first trimester.
<sup>c</sup> This is the number of high glycemic index foods consumed the day prior to the survey.

### Risk factors for gestational diabetes

Comparing the GDM present and absent groups, those who were normal weight (8.3% vs 7.8%) and underweight (6.3% vs 7.4%) per BMI categorization were statistical the same but the overweight group had significantly more GDM cases (35.4% vs 20.8). On the day prior to the survey, 28.6% of the GDM group consumed more than five high glycemic index foods while it was 18.5% in the non-GDM group (p=0.080). Presented in Table 3 is the Chi-square test comparing characteristics of cases and controls and the crude and adjusted binary logistic regressions showing the significant risk factors for GDM.

**Table 3.** Characteristic of cases and controls and unconditional binary logistic regression showing crude and adjusted odds ratio for gestational diabetes defined as 2-hour OGTT ≥8.5 mmol /l and/or the fasting plasma glucose ≥5.6 mmol/l

| Dichotomous risk factors | GDM<br>(n=70) | No GDM<br>(n=376) | *P-value* | Unconditional binary logistic regression |  |  |  |
| --- | --- | --- | --- | --- | --- | --- | --- |
|  |  |  |  | Univariate |  | Multivariate |  |
|  |  |  |  | Crude OR | 95% CI | Adj OR | 95% CI |
| ***Socio-demographic risks*** |  |  |  |  |  |  |  |
| Age >35 years | 16 (23.9) | 46 (12.0) | 0.019 | 2.29 | 1.21-4.36* | 4.06 | 0.58-8.73 |
| Unmarried | 13 (28.3) | 65 (27.3) | 0.859 | 1.05 | 0.52-2.12 | - | - |
| Rural residence | 14 (28.6) | 71 (29.1) | 0.941 | 0.98 | 0.49-1.92 | - | - |
| Low maternal education<sup>a</sup> | 11 (21.6) | 30 (12.0) | 0.077 | 2.01 | 0.931-4.33** | - | - |
| Low partner education<sup>a</sup> | 8 (16.7) | 16 (6.7) | 0.039 | 2.80 | 1.12-6.97* | 3.36 | 1.271- 8.89* |
| Primary facility | 16 (31.4) | 38 (15.1) | 0.009 | 2.56 | 1.29-5.08* | 4.95 | 1.87-3.76* |
| ***Anthropometry*** |  |  |  |  |  |  |  |
| Overweight/obese | 13 (20.0) | 39 (10.7) | 0.041 | 2.08 | 1.04-4.16* | 2.13 | 1.13-4.03* |
| Weight >90kg<sup>b</sup> | 4 (6.1) | 11 (3.0) | 0.182 | 2.08 | 0.64-6.75 | - | - |
| Height <150cm<sup>b</sup> | 7 (13.7) | 26 (10.4) | 0.466 | 1.37 | 0.56-3.35 | - | - |
| Weight gain >threshold<sup>c</sup> | 12 (24.0) | 51 (20.6) | 0.574 | 1.21 | 0.59-2.49 | - | - |
| MUAC >30cm<sup>c</sup> | 22 (34.9) | 80 (21.3) | 0.024 | 1.99 | 1.12-3.52* | 2.97 | 1.31-5.58* |
| ***Obstetric history*** |  |  |  |  |  |  |  |
| Parity >3 living children | 8 (12.9) | 22 (6.2) | 0.066 | 2.25 | 0.95-5.31** | 2.42 | 0.39-14.75 |
| Gravida >5 pregnancies | 5 (7.9) | 19 (5.0) | 0.365 | 1.63 | 0.58-4.53 | - | - |
| Prior macrosomia | 1 (16.7) | 8 (18.6) | 0.909 | 2.87 | 0.09-8.56 | - | - |
| Prior cesarean delivery | 10 (20.0) | 43 (16.3) | 0.539 | 1.28 | 0.59-2.75 | 4.01 | 1.09-14.76* |
| Prior neonatal death | 5 (10.2) | 21 (8.0) | 0.576 | 1.32 | 0.47-3.67 | 4.06 | 0.88-18.87** |
| Spontaneous abortion | 18 (50.0) | 62 (32.0) | 0.040 | 2.13 | 1.04-4.37* | 1.15 | 0.33-4.03 |
| Multiple pregnancies | 2 (4.0) | 7 (2.7) | 0.439 | 1.51 | 0.31-7.49 | - | - |
| ***Medical examinations*** |  |  |  |  |  |  |  |
| Diabetes in family | 5 (7.5) | 24 (6.3) | 0.787 | 1.20 | 0.44-3.26 | 1.50 | 0.31-7.31 |
| Family hypertension | 7 (13.7) | 22 (8.8) | 0.296 | 1.65 | 0.67-4.11 | 1.21 | 0.34-4.36 |
| Glycosuria<sup>d</sup> | 4 (5.5) | 11 (2.6) | 0.171 | 2.14 | 0.66-6.91 | 3.65 | 0.76-17.42 |
| Hypertension | 9 (17.6) | 47 (18.7) | 0.989 | 1.93 | 0.42-2.04 | 3.98 | 0.50-31.42 |
| Preeclampsia | 6 (9.1) | 6 (1.6) | 0.004 | 6.23 | 1.15-19.96* | - | - |
| Dyslipidemia<sup>e</sup> | 8 (15.7) | 63 (25.3) | 0.153 | 0.55 | 0.25-1.23 | 0.91 | 0.16-5.11 |
| HIV | 2 (5.1) | 2 (0.9) | 0.082 | 5.84 | 0.797-42.74 | - | - |
| **Nutrition** |  |  |  |  |  |  |  |
| Excess high GI foods<sup>f</sup> | 18 (28.6) | 56 (18.5) | 0.080 | 1.76 | 1.95-3.28* | 2.91 | 1.05-8.07* |
| Anemia<sup>g</sup> | 24 (60.0) | 130 (55.6) | 0.365 | 1.20 | 0.61-2.37 | - | - |

Model summary: number of observations = 358; Prob > $\chi^2$ = 0.0116; Log likelihood = -87.904; Pseudo $R^2$ = 0.2438.
BMI, body mass index; MUAC, mid-upper arm circumference.
Significant at \* $p < 0.05$ and \*\* $p < 0.10$
<sup>a</sup> Education up to the primary level was considered low.
<sup>b</sup> Weight and height were measured at ANC booking in the first trimester to assess BMI.
<sup>c</sup> MUAC was measured once in each trimester and weight was measured monthly. Per Institute of Medicine's recommendations on pregnancy weight gain based on BMI, a high GDM risk was considered if the woman's weight gain was above the threshold for her the BMI group.
<sup>d</sup> Dyslipidemia refers to total cholesterol >7.73 mmol/l, high-density lipoprotein cholesterol <1.34 mmol/l, low-density lipoprotein cholesterol >4.76 mmol/l and triglycerides >4.31 mmol/l.
<sup>e</sup> Glycosuria includes trace 1+ to 5+ dipstick glucose at any one time point in pregnancy.
<sup>f</sup> High caloric intake defined as excess intake of high glycemic index foods $\geq 5$ per day on the day prior to the survey.

In both the crude and adjusted models, spousal education below secondary level, ANC in a primary facility, adiposity (overweight/obesity, high MUAC) and intake of high glycemic index foods more than five times per day were significant risk factors for GDM. In addition, maternal age above 30 years, previous spontaneous abortion and preeclampsia in the current pregnancy were significant risk factors in the crude model. In the adjusted model, previous cesarean section was a significant independent risk factor (Table 3).

### Maternal and newborn outcomes

Records of 63 GDM cases and 340 controls were traced at birth. The maternal and newborn outcomes are presented in Tables 4 for continuous variables and Table 5 for categorical variables. Mean birth weight was 3.12 kg (SD=0.46) and was 0.26 kg higher among the cases (P=0.035). Similarly, estimated blood loss was 183.93 ml (SD=103.98) and was 50 ml higher among the cases (p=0.001). Deliveries of 21.5% (n=65) was by cesarean; it was 11.9% higher among the cases (P=0.049). No other significant variables were observed.

**Table 4.** Unpaired t-test comparing maternal and newborn birth outcomes for continuous variables among cases and controls

| Continuous outcomes | Mean (standard deviation) |  | P-value |
| --- | --- | --- | --- |
|  | GDM (n=63) | No GDM (n=340) |  |
| Blood loss (ml) | 228.42 (123.48) | 178.54 (105.89) | 0.010 |
| GA at birth (weeks) | 38.84 (2.19) | 38.80 (2.18) | 0.906 |
| Birth weight (kg) | 3.23 (0.49) | 3.06 (0.45) | 0.035 |
| Birth length (cm) | 49.59 (2.74) | 48.87 (3.34) | 0.212 |
| Head circumference (cm) | 34.38 (1.71) | 33.97 (2.04) | 0.253 |
| Ponderal index (g/cm<sup>3</sup>)<sup>a</sup> | 2.70 (0.49) | 2.69 (0.61) | 0.913 |
| Newborn glucose (mmol/l) | 3.924 (1.31) | 4.74 (0.90) | 0.193 |

GA; Gestational age.
<sup>a</sup> Ponderal Index calculated as newborn's weight (g)/ length (cm<sup>3</sup>) x 100

**Table 5.** Chi-square test comparing maternal and newborn birth outcomes for categorical variables among cases and controls

| Categorical outcomes | Number (percentage) |  | P-value<sup>a</sup> |
| --- | --- | --- | --- |
|  | GDM (n=63) | No GDM (n=340) |  |
| *Maternal outcomes* |  |  |  |
| Cesarean section | 16 (31.4) | 49 (19.5) | 0.049 |
| Type of cesarean |  |  | 0.125 |
| Emergency | 5 (35.7) | 25 (58.1) |  |
| Elective | 9 (64.3) | 18 (41.9) |  |
| Perineum |  |  | 0.369 |
| Intact | 25 (69.4) | 136 (75.2) |  |
| Episiotomy | 4 (11.2) | 25 (13.8) |  |
| Tear | 7 (19.4) | 20 (11.0) |  |
| Pre/eclampsia | 7 (13.7) | 22 (8.8) | 0.197 |
| Postpartum hemorrhage | 3 (7.9) | 7 (3.3) | 0.180 |
| Prolong/obstructed labor | 7 (15.6) | 26 (11.1) | 0.787 |
| *Newborn outcomes* |  |  |  |
| Sex |  |  | 0.332 |
| Male | 28 (60.9) | 116 (52.5) |  |
| Female | 18 (39.1) | 105 (47.5) |  |
| Gestational age at birth |  |  | 0.985 |
| Preterm <37 weeks | 3 (8.1) | 15 (8.7) |  |
| Term 37-42 weeks | 33 (89.2) | 154 (89.0) |  |
| Post-term >42 weeks | 1 (2.7) | 4 (2.3) |  |
| Growth for gestational age<sup>b</sup> |  |  | 0.870 |
| Small for gestational age | 2 (5.4) | 13 (6.2) |  |
| Marginal | 13 (35.1) | 82 (38.9) |  |
| Large for gestational age | 6 (16.2) | 40 (19.0) |  |
| Birth weight categories |  |  | 0.675 |
| Low birth weight | 2 (4.3) | 17 (7.6) |  |
| Normal | 43 (91.4) | 200 (89.3) |  |
| Macrosomia (≥4 kg) | 2 (4.3) | 7 (3.1) |  |
| Resuscitation | 15 (34.1) | 60 (27.0) | 0.219 |
| Intensive care | 1 (2.0) | 13 (5.3) | 0.289 |
| Perinatal death<sup>c</sup> | 1 (2.0) | 3 (1.2) | 0.525 |

<sup>a</sup> Postpartum hemorrhage is defined as estimated blood loss >500ml.
<sup>b</sup> Growth in relation to gestational age was estimated from Ponderal Index calculated as newborn weight (g)/ length (cm<sup>3</sup>) expressed as a percentage and categorized as small for gestational age (<2.0), marginal (2.0-2.5), normal (2.5-3.0.) and large for gestational age (≥3.0).
<sup>c</sup> Perinatal death includes both macerated and fresh cases

### Adverse pregnancy outcomes associated with gestational diabetes

As observed from the simple logistic regression, a unit increase in fasting plasma glucose and 2-hour OGTT was associated with a significant increase in birth weight and estimated blood loss (Table 6). From the multivariate binary logistic regression, GDM was associated with perineal tear whereas cesarean section was a significant outcome at the 10%-level. Regarding newborn outcomes, GDM was associated with birth asphyxia whereas large for gestational age and neonatal resuscitation were significant outcomes at the 10%-level (Table 7).

**Table 6.** Simple linear regression showing the coefficients for maternal and perinatal outcomes per unit rise in fasting plasma glucose and 2-hour OGTT

| Outcomes | Fasting plasma glucose |  |  | 2-hour OGTT values |  |  |
| --- | --- | --- | --- | --- | --- | --- |
|  | Coef. | 95% CI | P-value | Coef. | 95% CI | P-value |
| Cesarean section<sup>a</sup> | 0.185 | -0.087, 0.457 | 0.183 | 0.330 | -0.140, 0.801 | 0.168 |
| Episiotomy<sup>a</sup> | -0.235 | -0.601, 0.130 | 0.207 | -0.490 | -1.121, 0.140 | 0.127 |
| Perineal tear<sup>a</sup> | 0.204 | -0.168, 0.575 | 0.281 | 0.143 | -0.506, 0.793 | 0.664 |
| Preeclampsia<sup>a</sup> | 0.087 | -0.193, 0.368 | 0.541 | 0.149 | -0.339, 0.637 | 0.548 |
| Prolong labor | 0.028 | -0.098, 0.155 | 0.660 | 0.077 | -0.026, 0.122 | 0.200 |
| Est. blood loss | 0.196 | 0.087, -0.306 | 0.001* | 0.290 | 0.010-0.482 | 0.003* |
| Hemoglobin | 0.024 | -0.065, 0.114 | 0.592 | 0.043 | -0.105, 0.193 | 0.563 |
| Gestational age | 0.056 | -0.004, 0.116 | 0.067** | 0.034 | -0.072, 0.140 | 0.529 |
| Birth weight | 0.251 | 0.008, 0.494 | 0.043* | 0.562 | 0.141, 0.983 | 0.009** |
| Birth length | 0.001 | -0.034, 0.036 | 0.969 | 0.003 | -0.059, 0.065 | 0.923 |
| Head circumference | 0.056 | -0.001, 0.114 | 0.056** | 0.043 | -0.059, 0.147 | 0.405 |
| Apgar at 5min | -0.036 | -0.119, 0.064 | 0.558 | -0.064 | -0.236, 0.072 | 0.296 |
| Ponderal index<sup>b</sup> | 0.159 | -0.030, 0.349 | 0.100 | 0.273 | -0.060, 0.607 | 0.108 |
| Newborn glucose | 0.058 | -0.156, 0.273 | 0.583 | 0.029 | -0.420, 0.478 | 0.897 |
| Resuscitation<sup>a</sup> | 0.172 | -0.081, 0.426 | 0.181 | 0.272 | -0.142, 0.687 | 0.197 |
| Intensive care<sup>a</sup> | -0.286 | -0.0881, 0.307 | 0.343 | -0.734 | -1.757, 0.288 | 0.158 |
| Birth asphyxia<sup>a</sup> | 0.850 | -0.461, 2.163 | 0.203 | 0.457 | -1.792, 2.706 | 0.690 |
| Perinatal death<sup>ac</sup> | 0.719 | -0.353, 1.792 | 0.188 | 0.645 | -1.193, 2.484 | 0.490 |

Significant at \*p<0.05 and \*\*p<0.10
<sup>a</sup> Categorical variables while the rest are continuous variables.
<sup>b</sup> Ponderal Index derived from as fetal weight (g)/length (cm<sup>3</sup>)
<sup>c</sup> Perinatal death includes both macerated and fresh cases

**Table 7:** Adjusted odds for adverse pregnancy outcomes associated with gestational diabetes defined as 2-hour OGTT ≥8.5 mmol/l or the fasting plasma glucose ≥5.6 mmol/l

| Delivery outcomes | AOR | 95% CI | P-value |
| --- | --- | --- | --- |
| *Maternal outcomes* |  |  |  |
| Preeclampsia | 1.198 | 0.162-8.809 | 0.859 |
| Perineal tear | 2.909 | 1.081-5.566 | 0.043* |
| Cesarean section | 1.884 | 0.965-3.678 | 0.063** |
| Post-partum hemorrhage | 4.652 | 0.311-9.586 | 0.265 |
| *Newborn outcomes* |  |  |  |
| Preterm (<37 weeks) | 0.736 | 0.207-2.619 | 0.637 |
| Post-term (>42 weeks) | 1.227 | 0.133-11.330 | 0.856 |
| Small for gestational age<sup>a</sup> | 0.473 | 0.131-1.709 | 0.254 |
| Large for gestational age<sup>a</sup> | 2.661 | 0.863-5.048 | 0.082** |
| Resuscitation | 2.906 | 0.937-9.011 | 0.065** |
| Birth Asphyxia | 3.243 | 1.006-10.449 | 0.039* |
| Admission to NICU | 0.512 | 0.063-4.141 | 0.530 |
| Stillbirth | 2.382 | 0.211-26.824 | 0.482 |

Significant at \* $p<0.05$ and \*\* $p<0.10$
<sup>a</sup>Small for gestational age is birth weight <10<sup>th</sup> percentile for gestational age and large for gestational age large is large birth weight >90<sup>th</sup> percentile for gestational age

### Postpartum Glycemic status

At 12 weeks postpartum, 100 of the study participants who tested for GDM during pregnancy could be traced out of which 20 were diagnosed with GDM during pregnancy. Mean fasting plasma glucose of the GDM group in the second trimester of pregnancy was 5.70 mmol/l (SD=0.79). At 12 weeks postpartum, mean fasting plasma glucose of the GDM group had reduced to 4.39 (SD=0.83). Difference between the prenatal and postpartum fasting glucose was statistical (p=0.01). Out of the 20 GDM cases, 14 (70%) achieved euglycemia at 12 weeks postpartum while fasting plasma glucose of the remaining GDM cases 6 (30%) was persistently above 5.6 mmol/l at 12 weeks postpartum.

## Discussion

Findings show a GDM prevalence of 15.9%. The most important risk factors were excess intake of high glycemic index foods, adiposity and ANC in a primary facility. Key pregnancy outcomes were perineal trauma and birth asphyxia. Regarding post-delivery glycemia, 30% of the GDM cases remained hyperglycemic at 12 weeks postpartum. Evidence shows an increasing burden of GDM globally with majority of cases occurring in low- and middle-income countries [2]. Although GDM prevalence in Africa is relatively lower (9.5%), Africa records the highest proportion (73.7%) of all deaths attributable to diabetes but invests the least on diabetes healthcare [2].

Lower prevalence has been reported in African countries including Ghana (9.3%) [16], South African (9.1%) [21] and Nigeria (8.6%) [22] compared with developed regions like North America (12%) and Europe (14%) [2]. But Africa appears to be catching-up fast as pockets of high cases have recently been reported in Tanzania (19.5%) [23], South Africa (25.8%) [24] and Morocco (23.7%) [25]. However, diverse algorithms for testing and diagnosing pose challenges in comparing prevalence, risks, treatment effects, pregnancy outcomes, and harmonizing clinical practice [12]. In Tanzania for instance, a large variation was found between GDM prevalence per fasting glucose (18.3%) and 2-h OGTT (4.3%) [23].

Most studies on GDM in Africa are conducted in urban tertiary hospitals where prevalence is expectedly high. Interestingly, we noticed similar prevalence among secondary and primary facility users who are often peri-urban and rural dwellers. This could be explained in the context of rural dwellers having more children compared to urban dwellers [17] culminating in longer childbearing years and advance age at childbirth. In settings where fertility and abortion rates are high, reducing childbearing especially in older women is crucial. Also, primary healthcare users often have lower education and are more unlikely afford to healthcare. Adhering to treatment regimen and making healthy dietary choices could also be problematic. Besides, primary facilities are limited to providing basic EmONC services. Traveling longer distances for emergency EmONC increases risks for complications and bad obstetric outcomes. But undoubtfully, the nutrition transition poses risk for urban dwellers [2, 4, 7] who are significantly more obese (4.3% vs 13.5%), have higher triglycerides (35.0% vs 64.4%) [14] and double odds for obesity [16, 23].

A third [14] to half [7] of women with GDM have one or more risk factors. But some women develop GDM without any identifiable risk. Making diagnostic decision based on risk could miss half of all cases [14, 24] supporting the call to test all pregnant women [7]. Despite the diverse risk profile, adiposity and excess intake of high GI foods are striking. High carbohydrate intake has been associated with GDM in South Africa [26] where pregnant women believe that controlling for sugar and sweetener cravings during pregnancy is difficult to achieve [26]. Intensifying education on lifestyle modifications focusing on reduced intake of high GI foods, lower pregnancy weight gain in obese women and monitoring and intervening to ensure optimum glycemic control are crucial [7]. Midwives who are at forefront of ANC care need training on how to give woman-centered nutrition counseling that is acceptable and fits the socio-cultural context. Studies is needed to identify locally available and frequency consumed carbohydrate food sources that have low glycemic index.

Despite the diverse risks, effects of GDM on maternal and fetal outcomes are inconsistent [5, 8]. We noted that a unit rise in glucose significantly increased birth weight and blood loss. Postpartum hemorrhage is a common cause of maternal death in many low-resource settings and perineal trauma following vaginal delivery is a major risk. With 85% of annual global deliveries occurring in low- and middle-income countries coupled with the surge in GDM, efforts at reducing near-miss events and maternal mortality especially in weak health systems could be derailed if GDM detection, management and follow-up efforts are not intensified.

Few studies in Africa have assessed the effect of GDM on pregnancy outcomes. In Morocco for instance, birth weight was the only significant effect [25]. We noted an association of GDM with perineal tear, large-for-gestational age and birth asphyxia. Macrosomic babies have two-fold odds for cesarean birth and stillbirth in Ghana [27]. Women who deliver macrosomic babies per vaginal have higher risk for prolonged labour, perineal tear and shoulder dystocia [5, 8, 9]. The newborns whose lungs are not fully developed, are often hypoxic and require resuscitation and admission to NICU to establish extra-uterine breathing [27]. In primary facilities where EmONC services are unavailable, perinatal death becomes inevitable.

GDM is known to increase short- and long-term cardio-metabolic risk especially for type 2 diabetes [6, 7, 10]. A third of the GDM cases in our study was persistently hyperglycemic at 12 weeks postpartum. In Morocco where nutritional counselling and pharmacology intervention were given, 93.2% (n=110) attained glycemic control [25] but at 8 weeks postpartum, 50% of the intervention group had FPG values indicative of type 2 diabetes [25]. This situation is worrying and necessitates proper transition of the continuum of care from obstetricians and midwives to cadres of health professional who are versatile with diabetes care after the routine postpartum phase ends. By so doing, mother-offspring pairs who experience special events during pregnancy like GDM would be monitored, followed-up and supported to avert long-term complications.

There are some limitations worth highlighting. In determining BMI, first trimester weight was used instead of pre-pregnancy weight as that was the only possibility. But in the first trimester, substantial weight changes are not expected [20]. MUAC was useful in assessing maternal adiposity as a GDM risk and could be used when pregnancy is advanced, and BMI is no longer useful. As result of the global non-consensus on diagnostic thresholds for GDM, [7, 12], we restricted to using higher thresholds for 2-h OGTT (IADPSG/WHO: ≥8.5 mmol/l) and FPG (NICE: ≥5.6 mmol/l [11, 13] as the case definition. Higher diagnostic cut-offs are known to have higher sensitivity, better disease prediction [14], better within-patient correlation [5] and less likelihood for overdiagnosis [19]. Slightly different estimates could have been obtained if other diagnostic thresholds were used. The call for countries to develop national guidelines for screening and diagnosis considering resource availability [7] could facilitate context-specific diagnoses and care.

A key challenge in GDM testing is the ‘no show’ syndrome. While 23% of the pregnant women booked for GDM testing in Tanzania failed to attend their appointment [23], 45% was observed on our study. However, this did not affect the estimates as 50% attribution was accounted for in the design. Establishing postpartum contact with participants was difficult due to poor house addresses and relocations out of the study area. Out of the 70 GDM cases, only 20 (representing 28.6%) could be traced at 12 weeks postpartum. Hence the postpartum hyperglycemic estimates need to be interpreted in the context of the study population and cautiously generalized. Having observed elevated FPG among participants less than 20 years, and the underweight group, we cannot rule-out non-adherence to fasting requirements. The Indian strategy of performing 2-h OGTT irrespective of fasting state [7] could be considered in settings where adherence to overnight fasting is problematic. Encouraging partner support could improve diagnostic outcomes as we found higher partner education to reduce GDM risk significantly. A key strength is the follow-up to assess short-term birth outcomes and glycemic status. As previous obstetric outcomes were extracted from the ANC booklet, recall biases were minimized.

## Conclusion

Gestational diabetes was 5-27% depending on the test and diagnostic criteria. But prevalence based on 2-h OGTT was 9.0%, same as the 9.3% observed in the largest tertiary hospital in Ghana indicating a surge even in peri-urban and rural communities. This highlights the need to tackle rural-urban inequities in access to healthcare by equipping primary facilities with basic amenities to test, refer or manage cases. While we support testing all pregnant women using fasting plasma followed by 2-hour OGTT if available, consensus is needed on the diagnostic criteria which could be applicable for health systems in low resource settings who are the least prepared to handle implications of the surge in GDM prevalence. Physiologic interactions between fasting plasma glucose and 2-h postprandial glucose values during pregnancy should be further investigated. Considering the array of risk factors identified, concentrating on the modifiable risks through interventions that focus on lifestyle modification is key. Nutrition education tailored towards moderate intake of high glycemic index foods and dietary diversification, glycemic control and weight management in overweight/obese women is crucial. To avert long-term metabolic effects, there should be transition of care when the routine postpartum phase ends to ensure follow-up and monitoring of glycemic control among women who experience gestational diabetes.

## Abbreviations

ANC: Antenatal Care
FPG: Fasting Plasma Glucose
GDM: Gestational Diabetes Mellitus
GI: Glycemic index
IADPSG: International Association of Diabetes in Pregnancy Study Groups
MUAC: mid-upper arm circumference
NICE: National Institute for Health and Care Excellence
OGTT: Oral Glucose Tolerance Test
PI: Ponderal index
WHO: World Health Organization

## Acknowledgments

We thank the focal persons at the Volta regional hospital (Cynthia Agyapomaa), Ho municipal hospital (Rebecca Doku), Hohoe municipal hospital (Samuel Darrah), Margret Marquart Catholic Hospital (Patience Heh) and Jasikan District Hospital (Hope Akpeke) who coordinated the data collection. We also thank the study participants especially those who honored the follow-up visits.

